# Individual alpha power predicts language comprehension

**DOI:** 10.1101/2021.10.12.464065

**Authors:** P. Wang, Y. He, B. Maess, J. Yue, L. Chen, J. Brauer, A.D. Friederici, T.R. Knösche

## Abstract

Alpha power attenuation during cognitive task performing has been suggested to reflect a process of release of inhibition, increase of excitability, and thereby benefit the improvement of performance. Here, we hypothesized that changes in individual alpha power during the execution of a complex language comprehension task may correlate with the individual performance in that task. We tested this using magnetoencephalography (MEG) recorded during comprehension of German sentences of different syntactic complexity.

Results showed that neither the frequency nor the power of the spontaneous oscillatory activity at rest were associated with the individual performance. However, during the execution of a sentences processing task, the individual alpha power attenuation did correlate with individual language comprehension performance. Source reconstruction localized effects in temporal-parietal regions of both hemispheres. While the effect of increased task difficulty is localized in the right hemisphere, the difference in power attenuation between tasks of different complexity exhibiting a correlation with performance was localized in left temporal-parietal brain regions known to be associated with language processing.

From our results, we conclude that in-task attenuation of individual alpha power is related to the essential mechanisms of the underlying cognitive processes, rather than merely to general phenomena like attention or vigilance.

**Highlights:**

- Comprehension of structurally complexed embedded sentences is correlated with individual alpha power attenuation during task but not with alpha power at rest.
- These effects were localized in temporal-parietal brain regions known to be associated with language processing.

**Data availability statement:** Anonymized raw data will be made available upon request via email to the corresponding author provided the requesting researchers sign a formal data sharing agreement and cite this paper as origin of the data.

## 1. Introduction

The alpha band (8-12 Hz) typically forms the most stable and prominent peak in the EEG/MEG power spectrum (Berger, 1938; Schomer & Da Silva, 2012). These oscillations evidently play a major role in brain function at rest, as their power is attenuated in the task-relevant brain regions during various movement or cognitive tasks, including finger tapping, driving, arithmetic calculations, and sentence comprehension (Gastaldon et al., 2020; Klimesch et al., 1990; Magosso et al., 2019; Mann et al., 1996; Pfurtscheller, 1989; Wang et al., 2021). The power attenuation in the alpha band has also been shown to scale with task demand or engagement at the group level (Magosso et al., 2019; Wang et al., 2021). This phenomenon has been associated with cortical activation or release from inhibition due to the task (Klimesch, 2012; Pfurtscheller, 2003). It has also been shown, at the individual level, that the alpha power at rest or immediately before the task correlates with task performance (Jones et al., 2010; Van Dijk et al., 2008; van Ede et al., 2012), but evidence for such an individual relationship for the task-related power attenuation during task performance is scarce (Hilla et al., 2020). Therefore, the question remains whether the attenuation phenomenon is related to the essential mechanisms of the underlying cognitive processes, rather than merely to general phenomena like attention or vigilance.

Moreover, the characteristics of the alpha peak (e.g., frequency and power) are specific for each individual (Furman et al., 2018; Grabot & Kayser, 2020; Gulbinaite et al., 2017; Horschig et al., 2014; Katyal et al., 2019; Migliorati et al., 2020; Minami et al., 2020; Sadaghiani & Kleinschmidt, 2016; Smit et al., 2006). Hence, the somewhat non-univocal picture on the relationship between individual alpha power dynamics and task performance might also be rooted in the insufficient capture of the individual oscillations by using the classical broad frequency band (about 8-12 Hz).

In order to clarify the question if the individual alpha power attenuation is related to cognitive performance, we turn to the arguably most ‘human-like’ cognitive faculty, namely language. The ability to produce and understand language requires intense coordination of numerous cognition faculties, such as phonological perception, syntax processing, semantic association, working memory, attention, and motor control. Indeed, it has been shown that the individual alpha peak frequency as well as the alpha band power at rest are related to individual language abilities (Kwok et al., 2019; Rathee et al., 2020; Sklar et al., 1972). Recently, we have explored the alpha band power directly *during* a language task, and found an association between the alpha power attenuation and language task complexity (Wang et al., 2021). Consequently, we hypothesize that the difference in alpha power attenuation between language tasks of different complexity, gauged at the individual alpha peak frequency and in task-relevant brain regions, reflects the individual cognitive ability to handle the task, and hence might be associated with the observed performance.

In the present paper, we tested this hypothesis using the same data set as in our previously reported MEG study with healthy, native German adult speakers (Wang et al., 2021). On four days within a week, participants listened to complex German sentences, and answered probing questions regarding the thematic role assignment (“who is doing what to whom”). The sentences differed in their complexity by containing either single or double embedded relative clauses. Previously, we reported that the cortical (posterior superior temporal and adjacent parietal) alpha band power attenuation at the final embedding closure was significantly larger for double than for single embedded sentences (Wang et al., 2021). Here, rather than looking at the entire alpha band, we focus on the individual peak frequency obtained from the resting state. In particular, we examined the correlation between language performance and: (1) the individual peak frequency and its power at rest; (2) the power attenuation of the individual peak frequency during the experimental task at different syntactic positions in the sentences; (3) the spatial localization on the cortex for the observed correlation effects. The results confirm our hypothesis that the individual difference between the power attenuation for tasks of different complexities (double vs. single embedding sentence comprehension) correlates with the individual performance.

## 2. Materials and Methods

As we are reusing the data from our previous study, many of the methodological details are already described elsewhere (Wang et al., 2021). In the following, these aspects are just presented as brief summary.

### 2.1. Participants

Thirty right-handed native German speakers (fifteen females) were enrolled in this study (mean age: 27, range from 20 to 34). Their reading span was 3.7 ± 0.9 (mean ± SD). No neurological diseases or hearing impairments were reported. Participants were naïve to the purposes of the experiment and gave written informed consent prior to the experiment. The study was approved by the ethics committee of the University of Leipzig (number of this approval).

### 2.2. Stimulus material

Two types of German sentences with single and double hierarchical center embedding were presented (see Fig. S5). All sentences started with an introductory phrase followed by a relative clause initiated by a relative pronoun (e.g. dass / that). The beginning of each relative clause (brace) was labeled with *bxon* while the final verb of it was labeled with *bxoff* to indentify the same level of embedding. The place holder x represents the embedding level.

### 2.3. Experimental procedures

The experiment included four sessions carried out on four working days within one week. The stimuli were presented by the software ‘Presentation’ (www.neurobs.com). At each day, participants listened to 33 sentences of each sentence type (i.e. single and double embedded). Each participant received an individual randomization of all 264 sentences presented on the four days. None of the sentences was presented twice to the same participant. After each sentence, a content question was asked to test the understanding of the thematic role assignments. Each session comprised four blocks. Sentences were presented during the first three blocks. During the fourth block, resting-state was recorded for at least 10 minutes. Participants were asked to close their eyes and stay awake.

### 2.4. Behavioral data analysis

Behavioral performance was measured through the accuracy of the participants’ responses to the question task. In contrast to our previous study, the single valued total performance accuracy of each participant was estimated by a simple mean across the 4 experimental days.

### 2.5. MEG data acquisition and preprocessing

After preprocessing, the data of the first three blocks (task sections) were epoched of 0.5 s length starting with the event triggers at *b1on, b1off, b2on, b2off, b3on, and b3off* (on for embedding’s begin, and off for embedding’s closure). Data of the fourth block (rest section, total 12min) were epoched into 36 trails of 20s length each to get a frequency resolution of 0.05 Hz for the determination of the peak frequency. After artifact rejection, the median value of number of valid trials was 34, ranging from a minimum of 6 to a maximum of 35.

### 2.6. Estimation of individual spontaneous peak frequency in sensor space

The individual spontaneous peak frequencies were estimated from the preprocessed resting-state MEG recordings. For each subject on each day, the power spectrum density (PSD) for each sensor was estimated using the multi-taper method via the function *psd_multitaper* from the MNE-python v.0.16 (Gramfort et al., 2013) (using default setup, except normalization = ‘full’). The PSDs for each 20s-length trial were transformed to logarithmic scale (i.e., in dB) and then averaged cross trials and sensors. After the averaging, the 1/f pink noise background was estimated and subtracted from each average grand PSD. The maximal peaks between 7-29 Hz were identified as the individual peak frequency.

### 2.7. Estimation of individual peak frequency power during the task in source space

For source localization, we used individual single shell volume conductor models and source models constructed from the individual T1-weighted MR data. We utilized Freesurfer 6.0.0 to segment the inner skull as well as the cortical surface. Finally, the cortical surfaces were labeled according to Glasser et al. (2016). In this paper, we focused on the regions of interest (ROI), which showed significant alpha power difference for different sentence type at final closure of the embeddings *(b1off),* see Fig. 3A and Wang et al. (2021).

To estimate the alpha power in source space, we used the LCMV beamformer method (Van Veen et al., 1997) via the function *make_lcmv* by MNE-python v.0.16. The reconstructed current density was restricted to being perpendicular to the cortical surface. The noise covariance matrix was computed by the mean noise covariance from the empty room measurements obtained before and after each recording session. A data covariance matrix was computed separately for each day based on the whole sentence data. The PSD (in dB) of each source was estimated using the multi-taper method *(psd_multitaper*, data zero-padding to 2 s), separately for each subject, ROI, sentence type, and day as mean over all presented sentences. The 1/f pink noise background was estimated and subtracted from each average PSD separately. To estimate the spectral power of the individual peak frequency at task, we averaged the spectral power of the two frequency bins, which define the interval around the individual peak frequency at rest. Finally, relative power attenuation was calculated by normalizing to the respective *b1on* power value separately for each subject, ROI, sentence type, and day. Hence, all subsequently reported spectral power values in task conditions are relative power attenuations with respect to *b1on*.

For computing the individual frequency power in Yeo’s 17-networks (Yeo et al., 2011; Fig. 5A), we first morphed the individual source space to the fsaverage space via the source morph function in MNE-python v.0.16 (by setting fsaverage spacing “ico5”). The remaining steps were as same as for using the Glasser’s ROIs.

## 3. Results

### 3.1. The individual resting-state peak frequency is not correlated with the language performance

We first examined the relationship between the individual peak frequency during rest (averaged across four days) and the total performance accuracy. The individual peak frequency was estimated from the resting-state recordings at sensor level (for more details, see Materials and Methods section 2.6.). We first estimated it for each subject on each day (Fig. 1A) and then averaged it across the four days (Fig. 1C). The overall task performance was calculated for each subject by averaging the accuracy scores of the four experimental days, including, both, double and single embedded sentences. As expected, for most participants, their individual peak frequencies were located in the alpha frequency range (8-12 Hz; Fig. 1A,C). There were 6 participants, though, whose peak frequencies were estimated in the beta range (> 16 Hz). We found no statistical difference among the estimated individual peak frequencies of the four experimental days (Friedman’s test Q = 0.71, p = 0.87; Fig. 1B). We also found no clear association between the 4-day-average individual peak frequency and the total performance accuracy (Spearman’s correlation, r = 0.20, p = 0.29; Fig. 1D).

**Figure 1.**
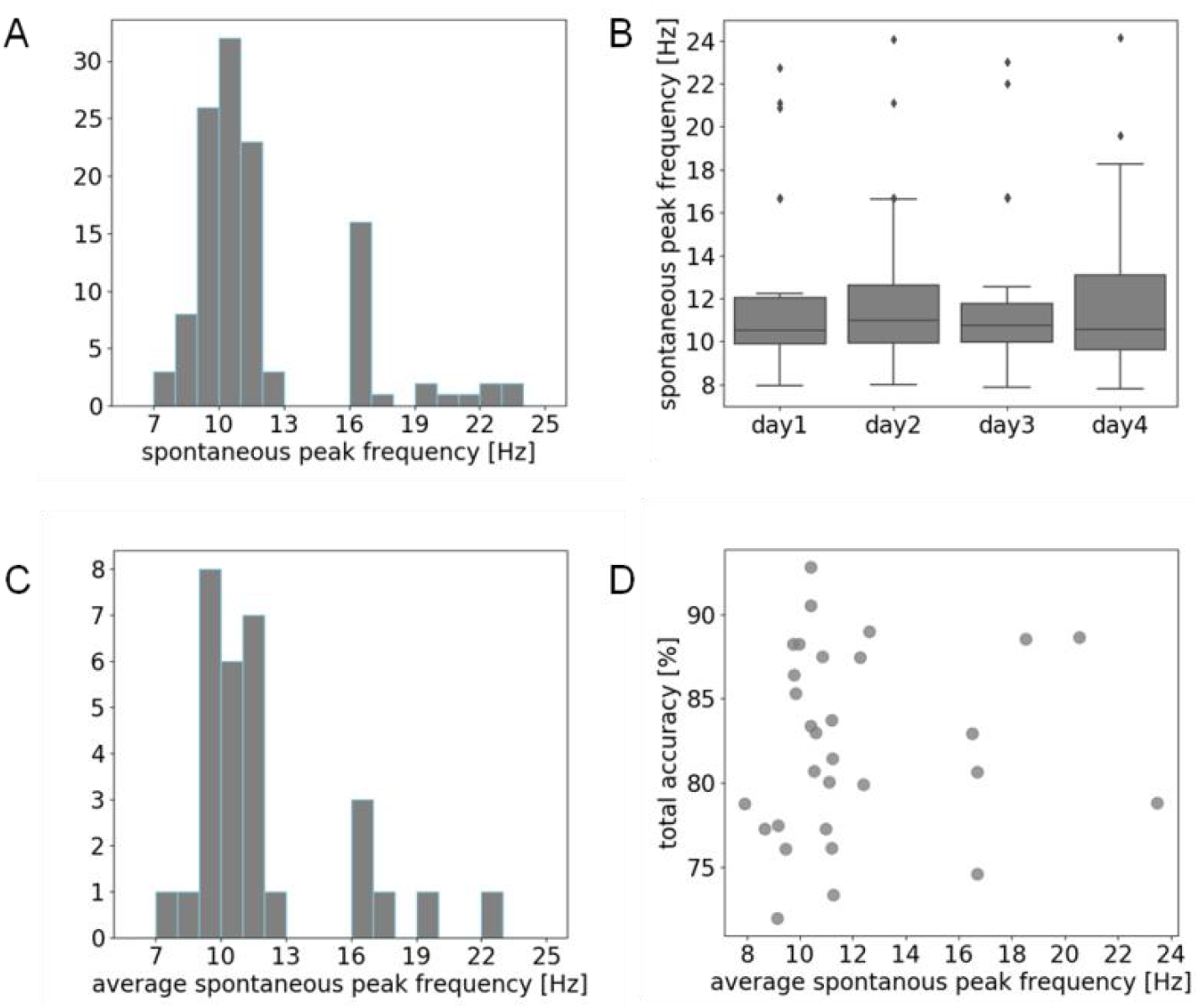
Relationship between the individual spontaneous peak frequency and the language task performance. (A) Histogram of the 120 spontaneous peak frequencies of the thirty participants on all four experimental days. (B) Boxplot of the participants’ individual spontaneous peak frequencies on each training day. The boxes show the interquartile ranges that stretch from the first quartile (25^th^ percentile) to the third quartile (75^th^ percentile) with the black line marking the median (50^th^ percentile). The maximal whisker range is 1.5 times the interquartile range. Note that the displayed whisker length depends on values within whisker range. Diamonds represent outliers, that is, values outside the whisker range. (C) Histogram of the thirty average spontaneous peak frequencies per person, averaged over the four experimental days. (D) Scatter plot showing the correlation between the total performance accuracy and the spontaneous peak frequency, both averaged over experimental days.

### 3.2. The power at the individual resting-state peak frequency is not associated with the language performance

Second, we examined the relationship between the power at the individual peak frequency during rest (average across four days) and the total performance accuracy. The power was also first estimated for each subject on each day (Fig. 2A) and then averaged across the four days (Fig. 2C). There were no detectable power differences between the four experimental days (Friedman’s test Q = 3.0, p = 0.39; Fig. 2B). We found no clear association between the (average) power and the performance accuracy (Spearman’s correlation, r = −0.09, p = 0.64; Fig. 2D).

**Figure 2.**
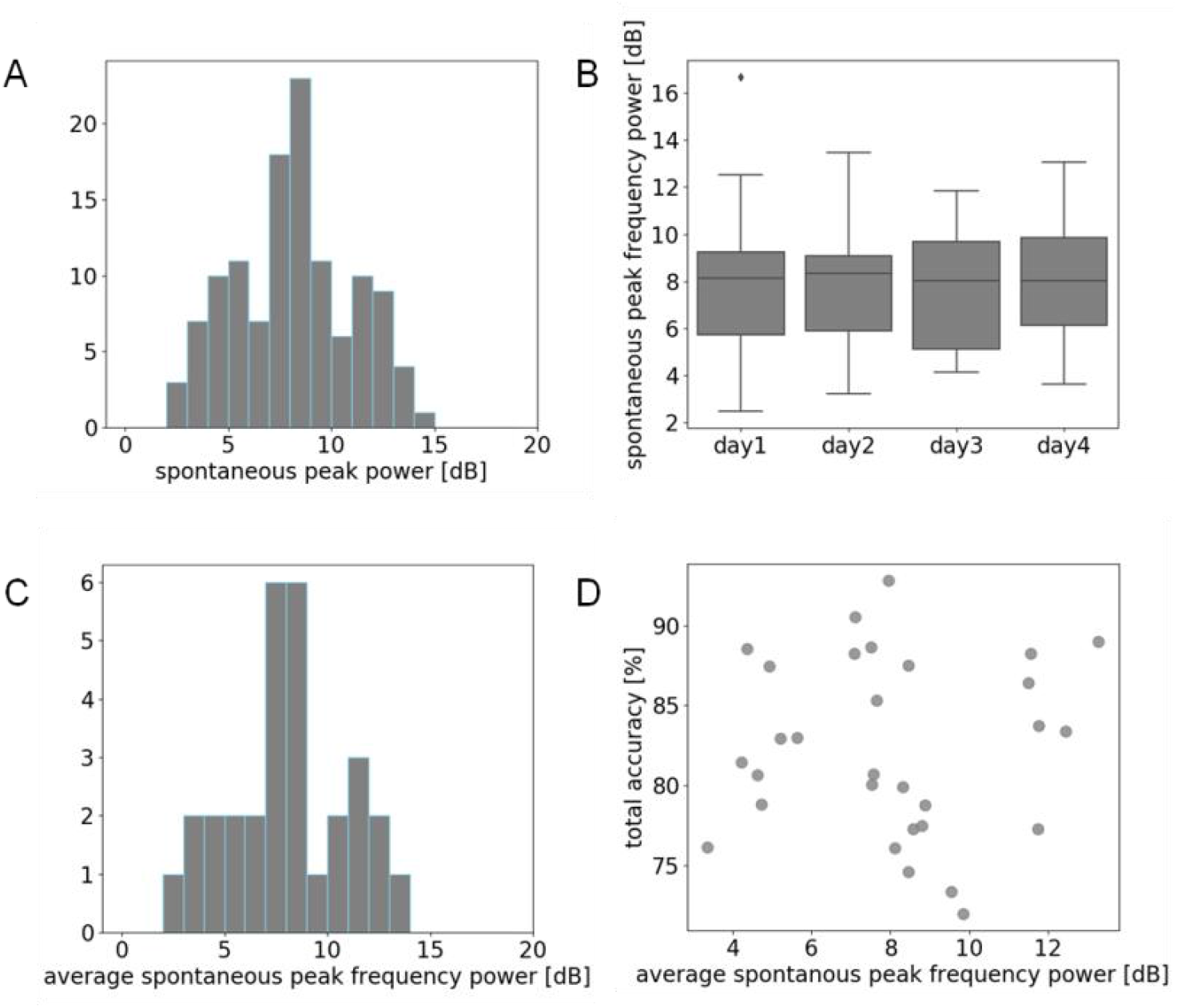
Relationship between the power of individual spontaneous peak frequency during rest and language task performance. (A) Histogram of the spontaneous peak frequency power of the thirty participants for all four experimental days (120 values). (B) Boxplot of participants’ individual spontaneous peak frequency power for each training day (for details, refer to Fig. 1). (C) Histogram of the spontaneous peak frequency power of the thirty participants, averaged over the four experimental days. (D) Scatter plot showing the correlation over participants between the total performance accuracy and the average spontaneous peak frequency power. Total performance accuracy was calculated by averaging the accuracy scores of the four experimental days, including, both, double and single embedded sentences. Average spontaneous peak frequency powers were averaged over experimental days (shown in C).

### 3.3. The power at the individual resting-state peak frequency during task was associated with the language performance

After examining the individual peak frequency and its power at rest, we examined the power at the same individual frequencies when participants were hearing the sentences, and tested its relationship with the language task performance at single subject level. We expected that the *difference* in power attenuation between the double embedded and single embedded sentences would reflect the subjects’ ability to process the sentences. To test this hypothesis, we focused on the brain regions (Fig. 3A), for which in our previous study (Wang et al., 2021) we found a significant difference in the alpha power (8-12 Hz) attenuation between the double and single embedding conditions at the final closure of the embedding structure *(b1off;* Fig. S5; for the meaning of the trigger point labels *b1on, b1off, b2on, b2off* see Materials and Methods section 2.2.). We first examined, whether by using the individual resting-state peak frequency, we can replicate our previous result obtained with the broad alpha band (sentence effect: general power attenuation difference between double and singled embedded sentences). In Fig. 3B we show the power decrease at the individual peak frequency at *b1off (b1off-b1on)* for each sentence type and each day. A non-parametric ANOVA (Wobbrock et al., 2011) for the factors sentence (double vs. single) and day (1 through 4) revealed only a significant main effect for the factor sentence (F = 26.10, p = 7.44E-7). Averaged across all four days, for both types of sentences, the alpha power decreases when sentence unfolds. The difference between the power attenuation between sentence types increased over the sentence and was most pronounced at *b1off* (Fig. 3C). The power attenuations at *b1off* for single and double embedded sentences were strongly correlated (Spearman’s correlation r = 0.84, p = 9.00E-9; Fig. 3D).

**Figure 3.**
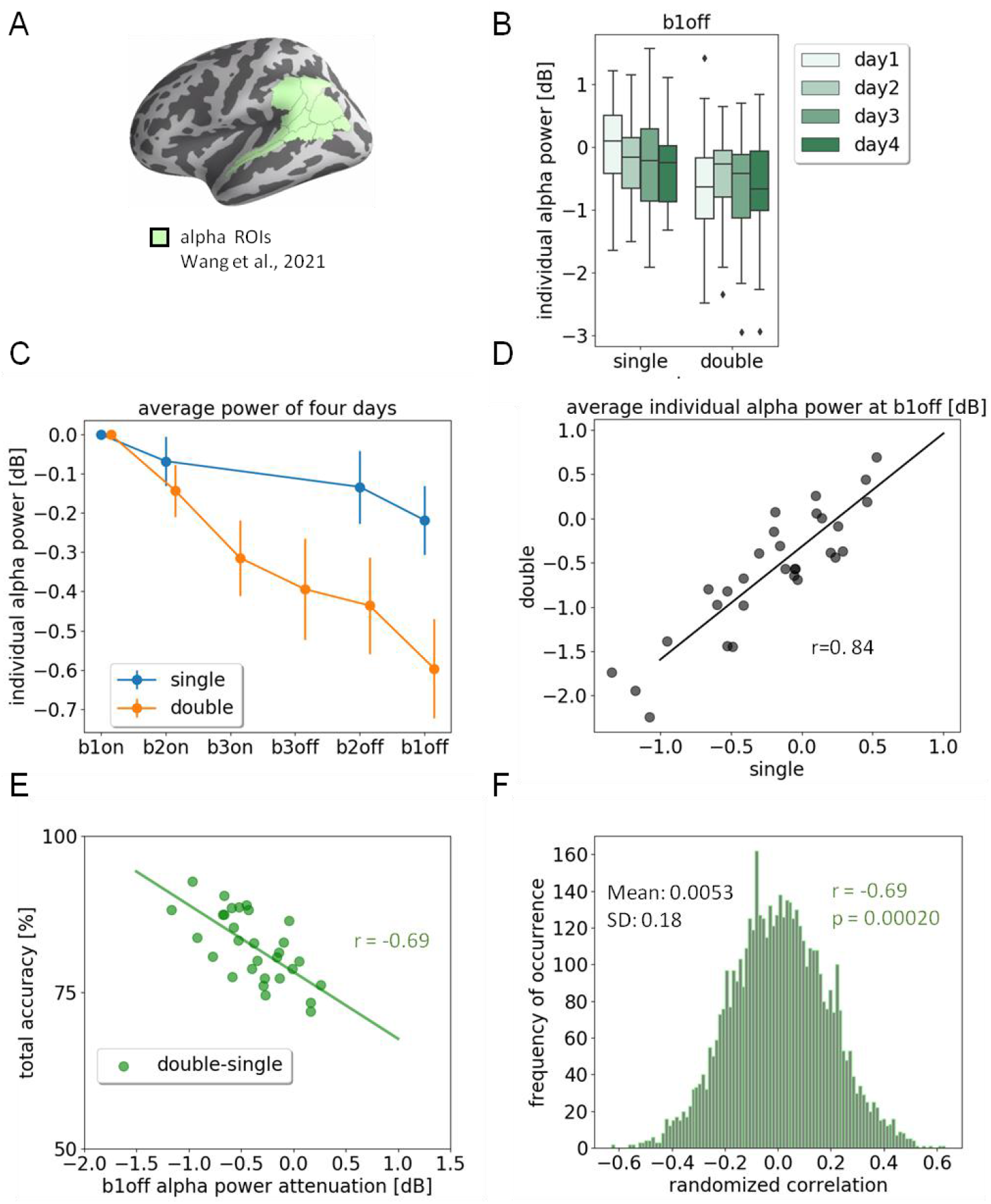
Relationship between the power attenuation at individual peak frequency during task and the language task performance. (A) Pre-selected ROIs that were reported to show significant power attenuation difference between double- and single embeddings at the final closure of the embedded structure in Wang et al., 2021. (B) Boxplot of the individual alpha power attenuation for each sentence type and each day. The individual alpha power attenuation was estimated by the power of the individual peak frequency at the final closure of the embedded structure (*b1off*) minus the power at the start of the embedded structure (*b1on*). (C) Four-days-average individual alpha power attenuation for each sentence type at different openings and closures of the embedded structure. Points show the mean value of the thirty participants, vertical lines show the standard error. The power of b1on (opening of the embeddings) was used as baseline for the other positions. (D) The individual alpha power attenuations at the final closures (*b1off*) for double and single embedded sentences were strongly associated (Spearman’s correlation, p = 9.00E-9). (E) The power attenuation differences between double and single embedded sentences at the individual alpha frequency at the final closure were associated with the individual performance accuracy (Spearman’s correlation, p = 2.63E-5). (F) Permutation-test of the association between the individual alpha power attenuation differences (double vs. single embedded sentence) at the final closure and the individual task performance. The power attenuations of the single as well as the double embedded sentences were permuted 5000 times. The mean of the randomized Spearman’s correlation was 0.0053 and the standard deviation was 0.18. The probability of appearance of a correlation value that less than r = −0.69 was 0.0002.

Most importantly, as hypothesized, at *b1off*, the alpha power attenuation difference between single and double embedded sentences was correlated with the individual task performance (Spearman’s correlation r = −0.64, p = 2.63E-5; Fig. 3E): participants who showed a larger power attenuation difference between the two sentence types at their individual peak frequency achieved better overall accuracy. Moreover, those participants, who showed a larger power attenuation for the double embedded sentences also performed better (Spearman’s r = −0.44, p = 0.015; see supplementary Fig. S1).

When using the classical frequency band (8-12 Hz, as reported in (Wang et al., 2021) instead of the individual peak frequency, the by-subject correlation between *b1off* power attenuation difference (double vs. single) and the performance reduced to r = −0.50 (Spearman’s correlation, p = 0.0049), and the correlation between the *b1off* power attenuation for double embedding and the performance dropped to r = −0.28 (Spearman’s correlation, p = 0.13; see supplementary Fig. S2).

### 3.4. Temporal exploration: association of alpha power attenuation and language performance at different embedding levels

Fig. 4A shows the Spearman’s correlation between the total performance accuracy and the individual peak frequency power attenuation (i.e., with respect to *b1on)* for both sentence types at different positions of the embedded structure. Fig. 4B shows, at individual peak frequency, the power attenuation *difference* between the two sentence types at equivalent positions: *innerOn/innerOff* and *outerOn/outerOff* (opening/closure of the innermost/outermost embedded structures). After FDR-correction, we found three significant correlations. In addition to the power attenuation at the final embedding closure for double embedded sentence and the power attenuation difference at the final embedding closure, which were already reported (Fig. 3E, & Fig. S1A), we found a performance correlation with the power attenuation difference at the first embedding closure *(innerOff;* Spearman’s r = −0.50, FDR p<0.05; Fig. 4B).

**Figure 4.**
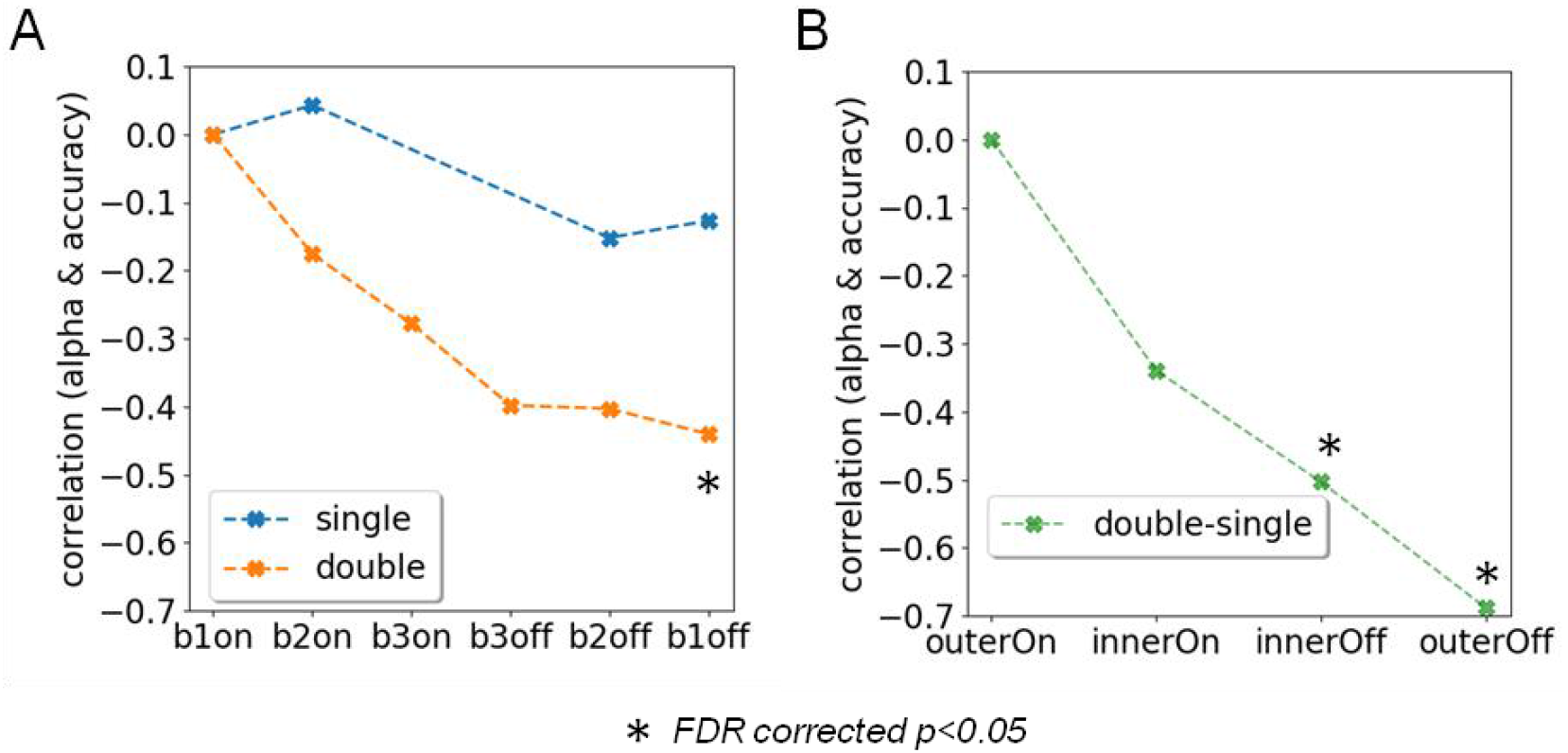
Association of the individual peak frequency power attenuation and the language performance at different positions of embedded structures. (A) Spearmans’ correlation between the individual peak frequency power attenuation (with respect to baseline b1on) for double and single embedded sentences and the total performance accuracy. (B) Spearmans’s correlation between the individual peak frequency power attenuation difference (double-single) and the total performance accuracy at different syntactically equivalent position. Position labels: outerOn: b1on for single and double; innerOn: b2on for single and b3on for double; innerOff: b2off for single and b3off for double; outerOff: b1off for single and double.

### 3.5. Spatial exploration: association of individual peak frequency power attenuation and language performance at the functional network level

We explored the correlation between the individual peak frequency power attenuation and the power attenuation difference (double vs. single embedding) of the individual peak frequency at *b1off* and the total performance at the cortical level. In order to achieve maximum specificity to functional networks, we used the Yeo’s 17-networks (Thomas Yeo et al., 2011) to define the brain ROIs (Fig. 5A). The power attenuation difference in the left temporal-parietal network (including the anterior and posterior superior temporal gyrus, Fig. 5A bright blue ROI) was significantly correlated with task performance (Spearman’s r = −0.62, FDR corrected p<0.05; Fig. 5B&C). Also, the power attenuation for double embedded sentence in the right temporal-parietal network (including anterior and posterior superior temporal gyrus, Fig. 5A bright blue ROI) was significantly correlated with the performance (Spearman’s r = −0.62, FDR corrected p<0.05; Fig. S3B&C).

**Figure 5.**
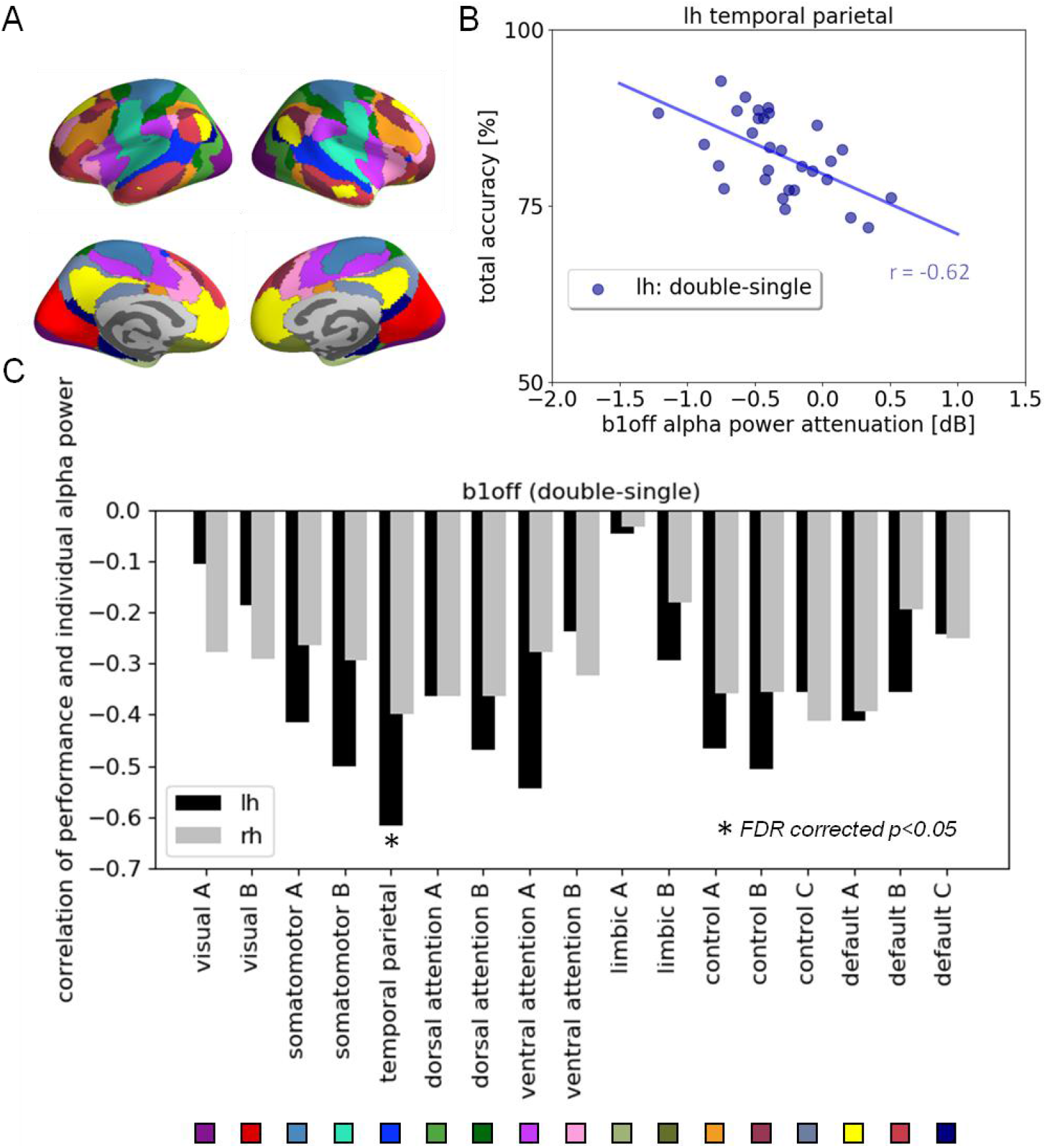
Association of the individual peak frequency power attenuation and the language performance at the functional networks level. (A) ROIs of Yeo’s 17-networks. The temporal parietal network is painted in bright blue. (B) Individual peak frequency power attenuation difference (double vs. single embeddings) at the final closure of the embedded structures (b1off) in the left temporal-parietal network (lh: double-single) was significantly associated with the performance (Spearmans’ r = – 0.62, FDR corrected p<0.05). (C) Spatial distribution of the Spearman’s correlation between individual peak frequency power attenuation difference (double vs. single embeddings) at the final closure of the embedded structures *(b1off)* and the individual performance accuracy on Yeo’s 17-networks.

Additional information about the spatial exploration of the correlation between the individual peak frequency power attenuation at the final closure of the embedded structure and the language performance based on the Glasser atlas (Glasser et al., 2016) can be found in Fig. S5. We obtained similar results as for the Yeo’s 17- networks: in the left anterior and posterior superior temporal gyri, the power attenuation difference between double and single embedded sentences was correlated with the language performance, while in the right middle and posterior superior temporal gyri, as well as inferior frontal gyrus, the power attenuation for double embedded sentences was correlated with the performance.

## 4. Discussion

In this study, we investigated the association between language comprehension performance and individual dominant oscillations under resting-state and in-task conditions. The individual peak frequency, as determined at sensor level at rest, was found in the alpha band for most participants. This individual peak frequency as well as the spectral power at that frequency at rest did not significantly change over the course of the experiment, nor did they correlate with the individual language performance.

Next, we turned to the MEG data acquired during task, by studying the spectral power attenuation at the individual peak frequency (as determined at rest). We tested our hypothesis that the individual power attenuation over the course of each sentence predicts the individual language comprehension performance. Indeed, we found correlations for, both, the power attenuation observed for the more complex sentences (double embedding) and the difference in power attenuation between the two complexity levels. While the same effects could be replicated when simply using the entire alpha band (8-12 Hz) instead of the individual peak frequency, the correlations were considerably weaker in that case. Interestingly, a spatial analysis based on a network atlas (Yeo’s 17-networks) suggested that the difference between the power attenuations is present in a temporal-parietal network in the left hemisphere, while the effect of the double embedding condition alone localizes in its right hemisphere homologue.

Hence, in summary, the following observations underscore the functional role of these individually dominant (mostly alpha band) oscillations in language processing: (i) individual performance was correlated with alpha power attenuation during task but not with alpha power at rest, and (ii) these effects were localized in temporal-parietal brain regions usually associated with language processing.

Cortical alpha activity has been proposed to play an important role in excitability regulation mechanisms underlying various human cognitive abilities, including memory, attention, perception, etc. Task irrelevant regions exhibit higher alpha power reflecting inhibition, while in task relevant regions alpha power is reduced, reflecting increased excitability (Foxe & Snyder, 2011; Jensen & Mazaheri, 2010; Klimesch, 2012; Klimesch et al., 2007).

At rest, such an increased excitability could reflect a general predisposition of a person towards cognitive performance. In contrast, if observed during task within specific task relevant brain areas, it would index processes related to the particular task or experimental situation. Alpha power at rest has been found to be positively correlated with task performance in cognitive control (Mahjoory et al., 2019) and episodic memory (Sargent et al., 2021) tasks, but negatively correlated with language skills (Kwok et al., 2019). This somewhat non-univocal picture suggests that for different experimental situations, different levels of pre-inhibition and disinhibition of cortical areas are beneficial. On the other hand, during the actual task and within the task relevant areas, we would expect a clear attenuation of alpha, which scales with the actual engagement with the task reflected by task performance. There are a number of previous studies showing general alpha attenuation during task within task relevant areas (Hilla et al., 2020; Magosso et al., 2019; Wang et al., 2021). These works show that the alpha oscillation power is related to the specific task demand. This is reflected in our results by the finding that the individual alpha power attenuation is stronger for the more complex sentences and increases with increasing cognitive (including working memory) load along the sentence. On the other hand, evidence for a correlation between alpha power modulation in specific task-relevant brain areas and individual task performance is scarce. By clearly demonstrating this in our study, we show that in-task alpha power attenuation actually reflects processes that are involved in successful task completion.

Along with the idea that the association of the alpha power and performance may indicate the task engagement of the brain regions, our results based on Yeo’s 17-networks suggest that the temporal-parietal network plays a crucial role in processing the embedding. We further cross-checked and refined this by exploring the association using the whole 360 ROI brain parcellation from the Glasser atlas and found a particularly high association between power attenuation difference (double vs. single) and performance in the superior temporal gyrus (Fig. S4. A). This is in alignment with the literature showing that the posterior superior temporal gyrus plays an important role in processing embedded structures (Friederici, 2011; Friederici et al., 2006; Kinno et al., 2008; Röder et al., 2002). Regarding the differential findings in the two homologue networks in the left and right hemispheres, we infer that even though the areas in the left hemisphere form the classical language network (Friederici, 2011; Hickok & Poeppel, 2004; Poeppel et al., 2012; Vigneau et al., 2006), homologue areas in the right hemisphere may increasingly engage when task demands are high as a result of enhanced working memory loads (Fridriksson & Morrow, 2005). This would explain why we find right hemispheric effects for the double embedded sentences, but not for the single embedded ones. Nevertheless, this notion alone would predict effects for both embedding levels in the left hemisphere as well. Instead, we only found an effect of the difference between these two levels, that is, on how much the alpha power attenuation increases when the task complexity rises. How could that be accounted for? One possible explanation is the following. Normal language processing only engages the left hemisphere language areas and leads there to an alpha power attenuation over the course of the sentence, which is strongly individually specific. Some persons have a large attenuation and others a smaller one, irrespective of the presented material. This causes a strong correlation between the alpha power attenuation between single and double embedded sentences (Fig. 3D). Then, there is an additional alpha power attenuation that is related to the cortical engagement with the task, and hence to sentence complexity (Fig. 3C) and task performance (Fig. 3E). While for each single condition (single and double embedding, respectively) this correlation is obscured by the large inter-individual fluctuation, in the difference between the two conditions this influence is cancelled out and the correlation with performance becomes significant. The right hemisphere areas, in contrast, are not so much involved in normal language processing and only step in to cope with the high task demand in the double embedding condition. Therefore, the alpha power attenuation is mainly task related and hence correlates with the task performance.

## 5. Conclusion

In summary, in this present paper, we used a language comprehension experiment to demonstrate that by manipulation of the task complexity, the alpha power attenuation in the task relevant brain regions, especially the individual alpha, reflect the increase of the cognitive load and predicted the individual performance outcome. Individual alpha power could be a useful biomarker to highlight the relevant brain regions and monitor the brain states during the cognitive processing.

## 6. Supplementary Figures

**Figure S1.**
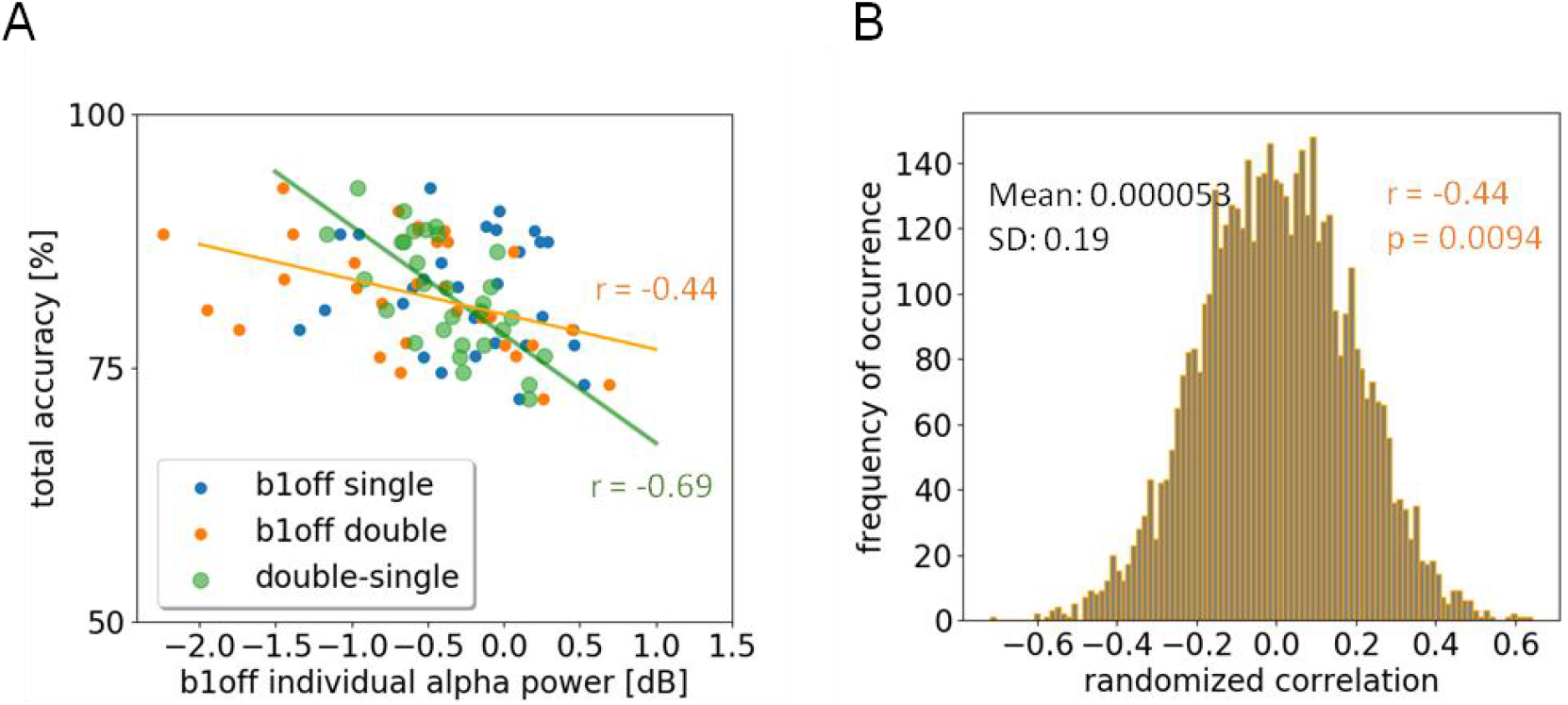
Association between the individual peak frequency power attenuation at the final closure of the embedded structure *(b1off)* and the total performance accuracy. (A) Individual peak frequency power attenuation for double embedded sentences was associated with the performance accuracy (Spearman’s r = −0.44, p = 0.015). (B) Permutation-test of the association between the individual peak frequency power attenuation (*b1off*) for double embedded sentences and the performance accuracy. The power attenuation values were permuted for 5000 times. The mean of the randomized Spearman’s correlation was 0.00 and the standard deviation was 0.19. The probability of appearance of a correlation value less than r = −0.44 was 0.0094.

**Figure S2.**
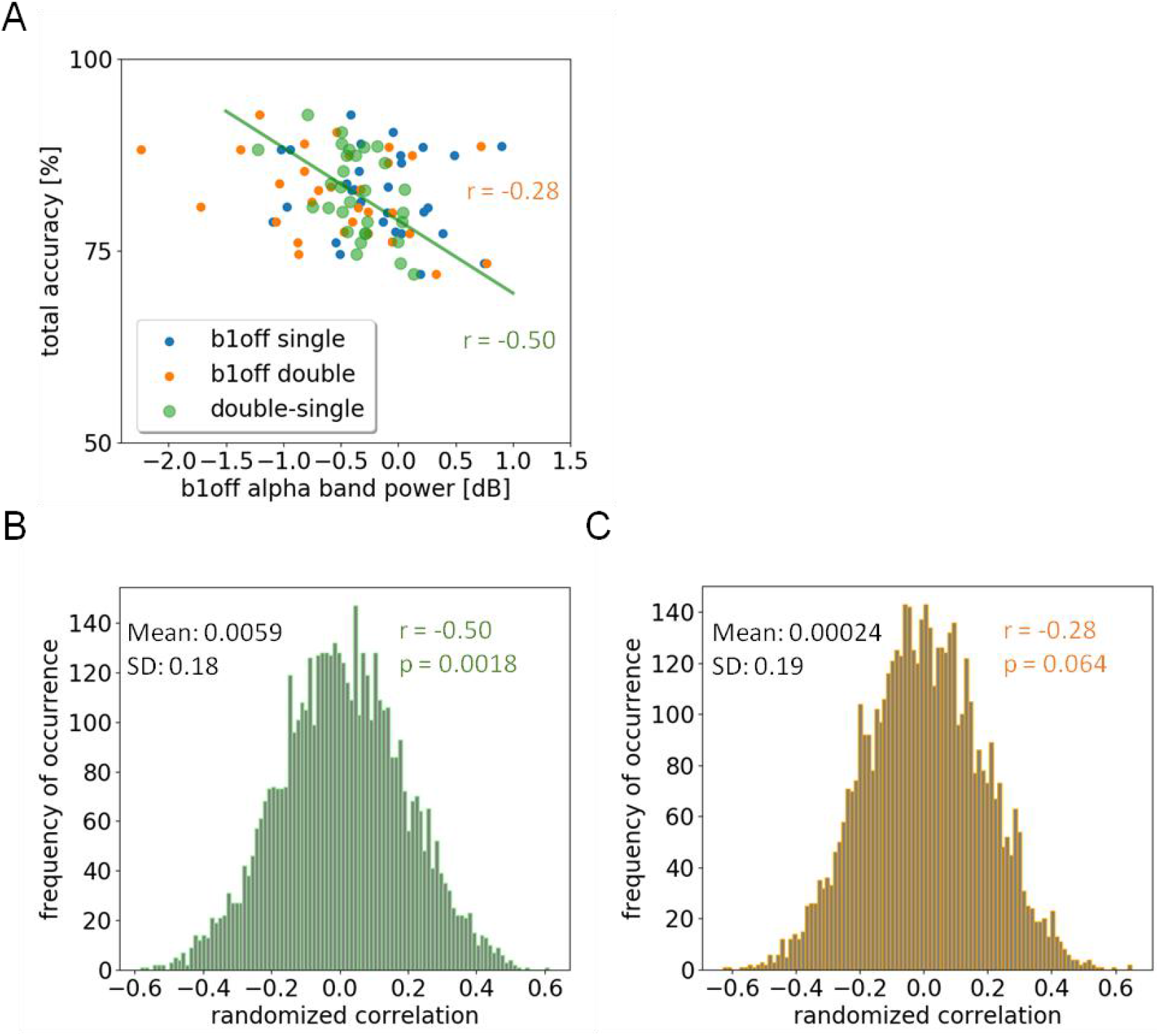
Association between the broad band alpha (8-12 Hz) power attenuation at the final closure of the embedded structure *(b1off)* and the total performance accuracy. (A) Broad band alpha power attenuation difference (double vs. single embeddings) associated with the total performance accuracy (Spearman’s r = −0.50, p = 0.0049). (B) Permutation-test of the association between the broad band alpha power attenuation difference (double vs. single embedded sentence) at the final closure of the embeddings and the individual task performance. The power attenuations of the single as well as the double embedded sentences were permuted for 5000 times. The mean of the randomized Spearman’s correlation was 0.0059 and the standard deviation was 0.18. The probability of appearance of a correlation value less than r = −0.50 was 0.0018. (C) Permutation-test of the association between the broad band alpha power attenuation (*b1off*) for double embedded sentences and the performance accuracy. The power attenuations were permuted for 5000 times. The mean of the randomized Spearman’s correlation was 0.00024 and the standard deviation was 0.19. The probability of appearance of a correlation value less than r = −0.28 was 0.064.

**Figure S3.**
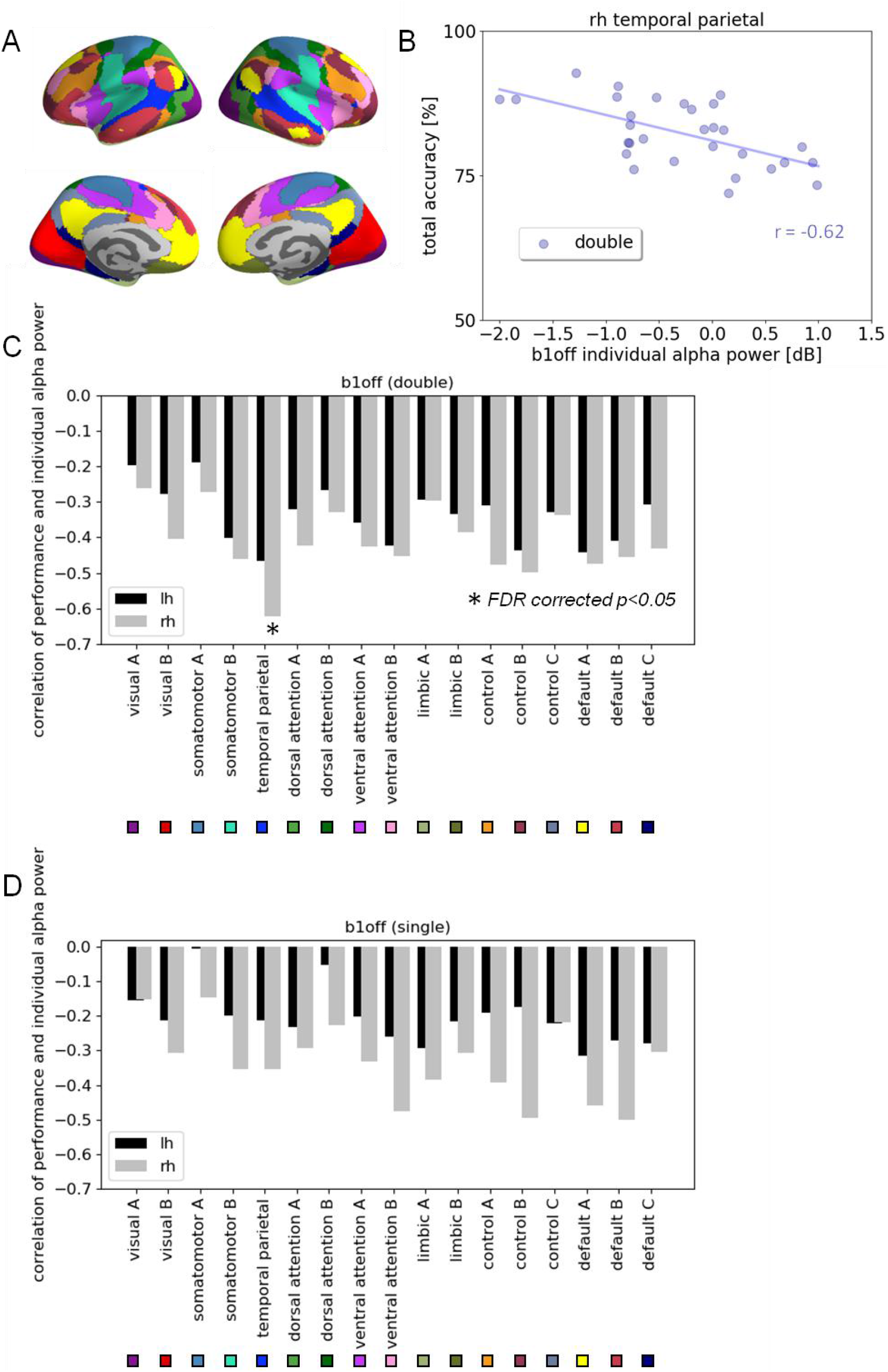
Association of the individual peak frequency power attenuation and the language performance at the functional networks. (A) ROIs of Yeo’s 17-networks. Temporal parietal networks are marked with bright blue. (B) Individual peak frequency power attenuation for double embedded sentences at the final closure of the embedded structures (*b1off*) on the right temporal-parietal network was significantly associated with the performance (Spearmans’ r = −0.62, FDR corrected p<0.05) (C)&(D) Spatial distribution of the Spearman’s correlation between individual peak frequency power attenuation of double (C) and single (D) embedded sentences at the final closure of the embedded structures (*b1off*) and the individual performance accuracy on Yeo’s 17-networks.

**Figure S4.**
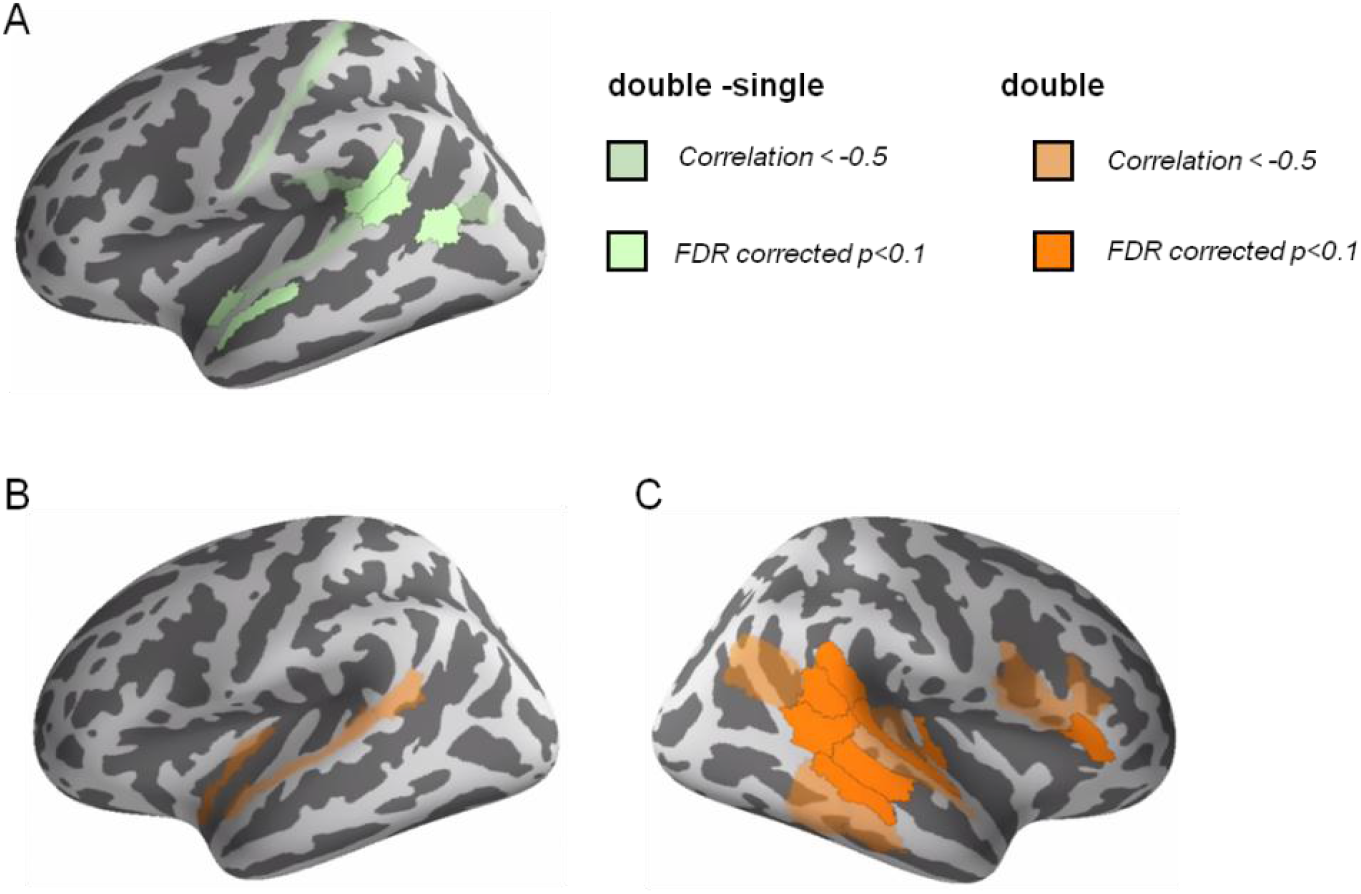
Association of the peak frequency power attenuation and the language performance with the Glasser atlas. Spearman’s correlation were calculated between the total performance accuracy and the (i) power attenuation difference (double vs. single embeddings) at the final closure of the embedded structure (b1off) as well as the power attenuation at the final closure for (ii) double and (iii) single embedded sentences for 360 ROIs. FDR-correction was based on all 1080 tests. Significant level is p<0.1 (A) ROIs that showed the association between power attenuation difference (douvle vs. Single embeddings) and the total performance accuracy. (B)&(C) ROIs that showed the association between power attenuation for double embedded sentences and the total performance accuracy.

**Figure S5.**
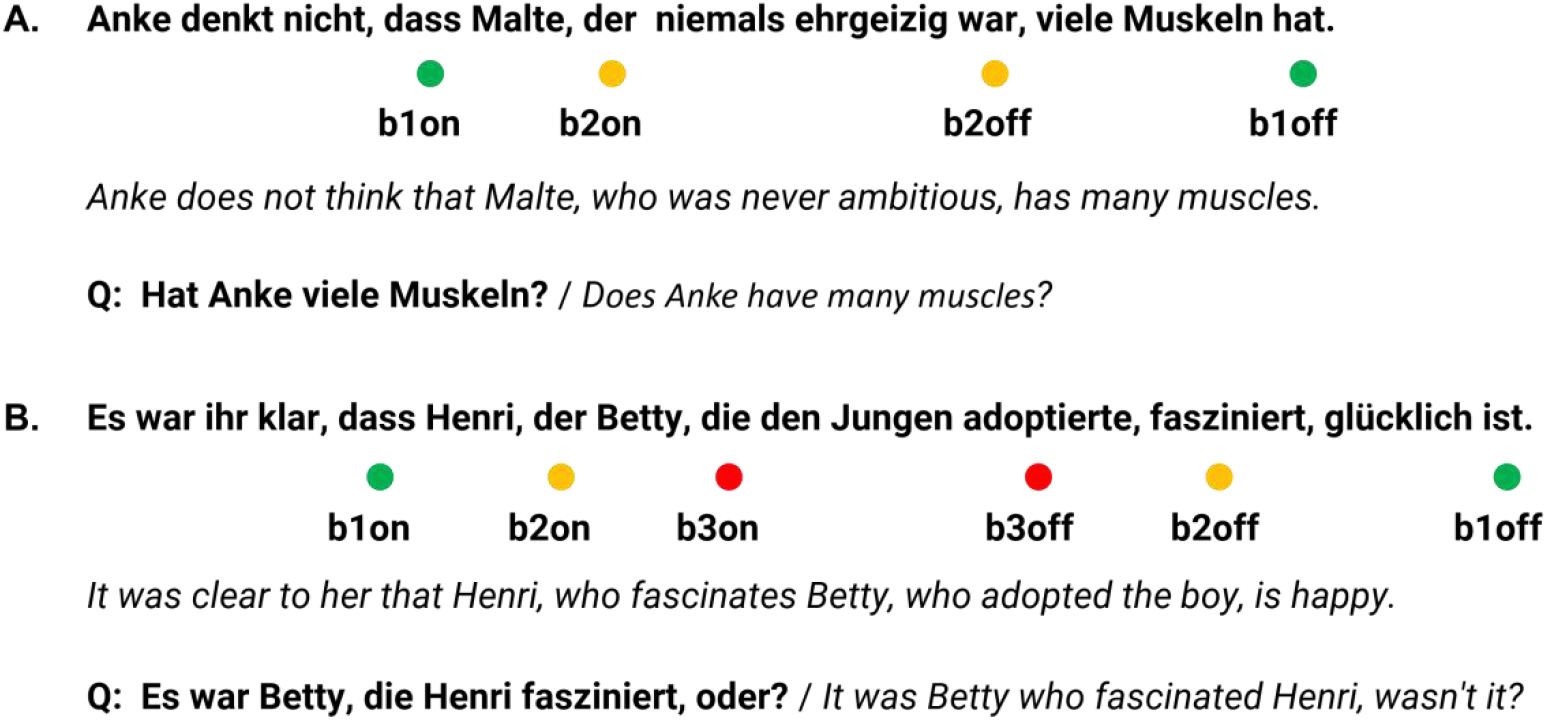
Examples of presented sentences in German with English translations in italics: (A) single; (B) double center embedding. We used the data segment starting at *b1on* as a reference to compute the relative alpha power attenuation for all other marked time points. The marker *b1on* represented the beginning of the relative clause containing all hierarchical embedding. Q: Probing questions for the presented examples.

## References

Berger, H. (1938). Über das Elektrenkephalogramm des Menschen. XIV. Archiv Für Psychiatrie Und Nerven kran kheiten.

Foxe, J. J., & Snyder, A. C. (2011). The role of alpha-band brain oscillations as a sensory suppression mechanism during selective attention. Frontiers in Psychology, 2(JUL), 1–13. https://doi.org/10.3389/fpsyg.2011.00154

Fridriksson, J., & Morrow, L. (2005). Cortical activation and language task difficulty in aphasia. Aphasiology, 19(3-5), 239–250.

Friederici, A. D. (2011). The brain basis of language processing: from structure to function. Physiological Reviews, 91(4), 1357–1392.

Friederici, A. D., Bahlmann, J., Heim, S., Schubotz, R. I., & Anwander, A. (2006). The brain differentiates human and non-human grammars: Functional localization and structural connectivity. Proceedings of the National Academy of Sciences of the United States of America, 103(7), 2458–2463. https://doi.org/10.1073/pnas.0509389103

Furman, A. J., Meeker, T. J., Rietschel, J. C., Yoo, S., Muthulingam, J., Prokhorenko, M., Keaser, M. L., Goodman, R. N., Mazaheri, A., & Seminowicz, D. A. (2018). Cerebral peak alpha frequency predicts individual differences in pain sensitivity. NeuroImage, 167, 203–210.

Gastaldon, S., Arcara, G., Navarrete, E., & Peressotti, F. (2020). Commonalities in alpha and beta neural desynchronizations during prediction in language comprehension and production. Cortex, 133, 328–345. https://doi.org/10.1016/j.cortex.2020.09.026

Glasser, M. F., Coalson, T. S., Robinson, E. C., Hacker, C. D., Harwell, J., Yacoub, E., Ugurbil, K., Andersson, J., Beckmann, C. F., & Jenkinson, M. (2016). A multi-modal parcellation of human cerebral cortex. Nature, 536(7615), 171–178.

Grabot, L., & Kayser, C. (2020). Alpha activity reflects the magnitude of an individual bias in human perception. Journal of Neuroscience, 40(17), 3443–3454.

Gramfort, A., Luessi, M., Larson, E., Engemann, D. A., Strohmeier, D., Brodbeck, C., Goj, R., Jas, M., Brooks, T., & Parkkonen, L. (2013). MEG and EEG data analysis with MNE-Python. Frontiers in Neuroscience, 7, 267.

Gulbinaite, R., van Viegen, T., Wieling, M., Cohen, M. X., & VanRullen, R. (2017). Individual alpha peak frequency predicts 10 Hz flicker effects on selective attention. Journal of Neuroscience, 37(42), 10173–10184.

Hickok, G., & Poeppel, D. (2004). Dorsal and ventral streams: a framework for understanding aspects of the functional anatomy of language. Cognition, 92(1-2), 67–99.

Hilla, Y., von Mankowski, J., Föcker, J., & Sauseng, P. (2020). Faster Visual Information Processing in Video Gamers Is Associated With EEG Alpha Amplitude Modulation. Frontiers in Psychology, 11, 33–33.

Horschig, J. M., Jensen, O., van Schouwenburg, M. R., Cools, R., & Bonnefond, M. (2014). Alpha activity reflects individual abilities to adapt to the environment. NeuroImage, 89, 235–243.

Jensen, O., & Mazaheri, A. (2010). Shaping functional architecture by oscillatory alpha activity: gating by inhibition. Frontiers in Human Neuroscience, 4, 186.

Jones, S. R., Kerr, C. E., Wan, Q., Pritchett, D. L., Hämäläinen, M., & Moore, C. I. (2010). Cued spatial attention drives functionally relevant modulation of the mu rhythm in primary somatosensory cortex. Journal of Neuroscience, 30(41), 13760–13765.

Katyal, S., He, S., He, B., & Engel, S. A. (2019). Frequency of alpha oscillation predicts individual differences in perceptual stability during binocular rivalry. Human Brain Mapping, 40(8), 2422–2433.

Kinno, R., Kawamura, M., Shioda, S., & Sakai, K. L. (2008). Neural correlates of noncanonical syntactic processing revealed by a picture-sentence matching task. Human Brain Mapping, 29(9), 1015–1027.

Klimesch, W. (2012). Alpha-band oscillations, attention, and controlled access to stored information. Trends in Cognitive Sciences, 16(12), 606–617. https://doi.org/10.1016/j.tics.2012.10.007

Klimesch, W., Pfurtscheller, G., Mohl, W., & Schimke, H. (1990). Event-related desynchronization, ERD-mapping and hemispheric differences for words and numbers. International Journal of Psychophysiology, 8(3), 297–308.

Klimesch, W., Sauseng, P., & Hanslmayr, S. (2007). EEG alpha oscillations: the inhibition–timing hypothesis. Brain Research Reviews, 53(1), 63–88.

Kwok, E. Y. L., Cardy, J. O., Allman, B. L., Allen, P., & Herrmann, B. (2019). Dynamics of spontaneous alpha activity correlate with language ability in young children. Behavioural Brain Research, 359, 56–65. https://doi.org/10.1016/j.bbr.2018.10.024

Magosso, E., De Crescenzio, F., Ricci, G., Piastra, S., & Ursino, M. (2019). EEG alpha power is modulated by attentional changes during cognitive tasks and virtual reality immersion. Computational Intelligence and Neuroscience, 2019. https://doi.org/10.1155/2019/7051079

Mahjoory, K., Cesnaite, E., Hohlefeld, F. U., Villringer, A., & Nikulin, V. V. (2019). Power and temporal dynamics of alpha oscillations at rest differentiate cognitive performance involving sustained and phasic cognitive control. Neuroimage, 188, 135–144.

Mann, C. A., Sterman, M. B., & Kaiser, D. A. (1996). Suppression of EEG rhythmic frequencies during somato-motor and visuo-motor behavior. International Journal of Psychophysiology, 23(1–2), 1–7.

Migliorati, D., Zappasodi, F., Perrucci, M. G., Donno, B., Northoff, G., Romei, V., & Costantini, M. (2020). Individual alpha frequency predicts perceived visuotactile simultaneity. Journal of Cognitive Neuroscience, 32(1), 1–11.

Minami, S., Oishi, H., Takemura, H., & Amano, K. (2020). Inter-individual differences in occipital alpha oscillations correlate with white matter tissue properties of the optic radiation. Eneuro, 7(2).

Pfurtscheller, G. (1989). Functional Topography During Sensorimotor Activation Studied with Event-Related Desynchronization Mapping. Journal of Clinical Neurophysiology, 6(1), 75–84. https://doi.org/10.1097/00004691-198901000-00003

Pfurtscheller, G. (2003). Induced Oscillations in the Alpha Band: Functional Meaning. Epilepsia, 44(12 SUPPL.), 2–8. https://doi.org/10.1111/j.0013-9580.2003.12001.x

Poeppel, D., Emmorey, K., Hickok, G., & Pylkkänen, L. (2012). Towards a new neurobiology of language. Journal of Neuroscience, 32(41), 14125–14131.

Rathee, S., Bhatia, D., Punia, V., & Singh, R. (2020). Peak Alpha Frequency in Relation to Cognitive Performance. Journal of Neurosciences in Rural Practice, 11(3), 416–419. https://doi.org/10.1055/s-0040-1712585

Röder, B., Stock, O., Neville, H., Bien, S., & Rösler, F. (2002). Brain activation modulated by the comprehension of normal and pseudo-word sentences of different processing demands: a functional magnetic resonance imaging study. Neuroimage, 15(4), 1003–1014.

Sadaghiani, S., & Kleinschmidt, A. (2016). Brain networks and α-oscillations: structural and functional foundations of cognitive control. Trends in Cognitive Sciences, 20(11), 805–817.

Sargent, K., Chavez-Baldini, U., Master, S. L., Verweij, K. J. H., Lok, A., Sutterland, A. L., Vulink, N. C., Denys, D., Smit, D. J. A., & Nieman, D. H. (2021). Resting-state brain oscillations predict cognitive function in psychiatric disorders: A transdiagnostic machine learning approach. NeuroImage: Clinical, 30, 102617.

Schomer, D. L., & Da Silva, F. L. (2012). Niedermeyer’s electroencephalography: basic principles, clinical applications, and related fields. Lippincott Williams & Wilkins.

Sklar, B., Hanley, J., & Simmons, W. W. (1972). An EEG experiment aimed toward identifying dyslexic children. Nature, 240(5381), 414–416.

Smit, C. M., Wright, M. J., Hansell, N. K., Geffen, G. M., & Martin, N. G. (2006). Genetic variation of individual alpha frequency (IAF) and alpha power in a large adolescent twin sample. International Journal of Psychophysiology, 61(2), 235–243.

Thomas Yeo, B. T., Krienen, F. M., Sepulcre, J., Sabuncu, M. R., Lashkari, D., Hollinshead, M., Roffman, J. L., Smoller, J. W., Zöllei, L., & Polimeni, J. R. (2011). The organization of the human cerebral cortex estimated by intrinsic functional connectivity. Journal of Neurophysiology, 106(3), 1125–1165.

Van Dijk, H., Schoffelen, J.-M., Oostenveld, R., & Jensen, O. (2008). Prestimulus oscillatory activity in the alpha band predicts visual discrimination ability. Journal of Neuroscience, 28(8), 1816–1823.

van Ede, F., Köster, M., & Maris, E. (2012). Beyond establishing involvement: quantifying the contribution of anticipatory α-and β-band suppression to perceptual improvement with attention. Journal of Neurophysiology, 108(9), 2352–2362.

Van Veen, B. D., Van Drongelen, W., Yuchtman, M., & Suzuki, A. (1997). Localization of brain electrical activity via linearly constrained minimum variance spatial filtering. IEEE Transactions on Biomedical Engineering, 44(9), 867–880.

Vigneau, M., Beaucousin, V., Hervé, P.-Y., Duffau, H., Crivello, F., Houde, O., Mazoyer, B., & Tzourio-Mazoyer, N. (2006). Meta-analyzing left hemisphere language areas: phonology, semantics, and sentence processing. Neuroimage, 30(4), 1414–1432.

Wang, P., Knösche, T. R., Friederici, A. D., Maess, B., Chen, L., & Brauer, J. (2021). Functional brain plasticity during L1 training on complex sentences: Changes in gamma-band oscillatory activity. Human Brain Mapping, March, 1–13. https://doi.org/10.1002/hbm.25470

Wobbrock, J. O., Findlater, L., Gergle, D., & Higgins, J. J. (2011). The aligned rank transform for nonparametric factorial analyses using only anova procedures. Proceedings of the SIGCHI Conference on Human Factors in Computing Systems, 143–146.

